# Prion-like α-synuclein pathology in the brains of infants: Krabbe disease as a novel seed-competent α-synucleinopathy

**DOI:** 10.1101/2021.10.12.463948

**Authors:** Christopher Hatton, Simona S. Ghanem, David J. Koss, Ilham Y. Abdi, Elizabeth Gibbons, Rita Guerreiro, Jose Bras, International DLB Genetics Consortium, Lauren Walker, Ellen Gelpi, Wendy Heywood, Tiago F. Outeiro, Johannes Attems, Bobby McFarland, Rob Forsyth, Omar M. El-Agnaf, Daniel Erskine

**Author notes:** Corresponding author: Dr Daniel Erskine. Members listed in Supplementary Author List (International DLB Genetics Consortium).

## Abstract

Krabbe disease (KD) is an infantile neurodegenerative disorder resulting from pathogenic variants in the *GALC* gene which causes accumulation of the toxic sphingolipid psychosine. *GALC* variants are associated with increased risk of Lewy body diseases (LBD), an umbrella term for age-associated neurodegenerative diseases in which the protein α-synuclein aggregates into Lewy bodies. To explore whether α-synuclein in KD has pathological similarities to that in LBD, we compared *post-mortem* KD tissue to that of infant control cases and identified alterations to α-synuclein localisation and expression of modifications associated with LBD. To determine whether α-synuclein in KD displayed pathogenic properties associated with LBD we evaluated its seeding capacity using the real-time quaking-induced conversion assay. Strikingly, seeded aggregation of α-synuclein resulted in the formation of fibrillar aggregates similar to those observed in LBD, confirming the prion-like capacity of KD-derived α-synuclein. These observations constitute the first report of prion-like α-synuclein in the brain tissue of infants and challenge the putative view that α-synuclein pathology is merely an age-associated phenomenon, instead suggesting it can result from alterations to biological processes such as sphingolipid homeostasis. Our findings have important implications for understanding the mechanisms underlying Lewy body formation in LBD.

## Introduction

Krabbe disease (KD) is a rare autosomal recessive neurodegenerative disorder characterised by disrupted sphingolipid homeostasis. It is caused by mutations in the *GALC* gene, that encodes the lysosomal enzyme β-galactocerebrosidase responsible for degrading galactosylceramide [1]. Loss of β-galactocerebrosidase activity leads to the accumulation of the cytotoxic sphingolipid psychosine/galactosylsphingosine. Accumulation of psychosine may promote demyelination by promoting the formation of characteristic macrophages termed globoid cells, or by direct toxicity to oligodendrocytes [2]. In contrast to the infantile presentation of KD, Lewy body diseases (LBD), such as Parkinson’s disease (PD) and dementia with Lewy bodies (DLB), are the second most common age-associated neurodegenerative disorder. The aggregation of the protein α-synuclein into insoluble assemblies is thought to be central to the pathogenic mechanism causing these disorders, and is a major target for candidate therapeutics [3].

Although KD and LBD occur at the opposite ends of the age spectrum, there are compelling reasons to suggest they share some mechanistic overlap. Firstly, *GALC* polymorphisms are associated with idiopathic PD at the genome-wide level [4]. Secondly, psychosine, which accumulates throughout the KD brain, promotes the aggregation of α-synuclein into forms associated with LBD both *in vitro* and *in vivo* [5]. Recent studies limited to *post-mortem* analysis of frontal cortex tissue from three cases have also identified neuronal α-synuclein deposits in *post-mortem* Krabbe disease tissues [6], but these have employed antibodies recognising both physiological and pathological forms of α-synuclein and, thus, the extent to which α-synuclein in KD recapitulates key pathogenic features of LBD is not known.

There is growing interest in the LBD field regarding the importance of lipids in α-synuclein aggregation, based in part on recent observations that Lewy bodies consist of considerable amounts of lipid-bound organelles as well as α-synuclein protein [7, 8]. Therefore, we sought to investigate the extent to which α-synuclein in KD, a disorder of sphingolipid metabolism, recapitulates pathogenic features observed in LBD to better understand the potential mechanistic overlap between these disorders. We conducted histological and molecular analysis of autoptic KD brains and compared these to infant control tissue, reporting important similarities between α-synuclein in KD and LBD that constitute the first evidence, to our knowledge, of prion-like α-synuclein in the brains of infants.

## Results and discussion

### Aggregated α-synuclein in the KD brain

Initial neuropathological survey of *post-mortem* KD brain tissue with α-synuclein antibodies did not identify any Lewy bodies or swollen neurites characteristic of LBD in any of the cases. We identified physiological α-synuclein to be variably present within white matter globoid cells, particularly in the medulla oblongata, crus cerebri and cerebral cortex, as visualised by the pan-α-synuclein antibody, Syn1 (Figure 1, A; Supplementary Figure 1-2). Occasional globoid cells were also immunoreactive for the autophagic marker p62/SQSTM1, which labels cellular waste and thus its accumulation may indicate impaired autophagic processes (Figure 1, B-C). However, globoid cells were not recognised by antibodies against LBD-associated forms of α-synuclein such as aggregate-specific antibodies (Syn-O2 and 5G4), or α-synuclein phosphorylated at serine 129 (pS129).

**Figure 1:**
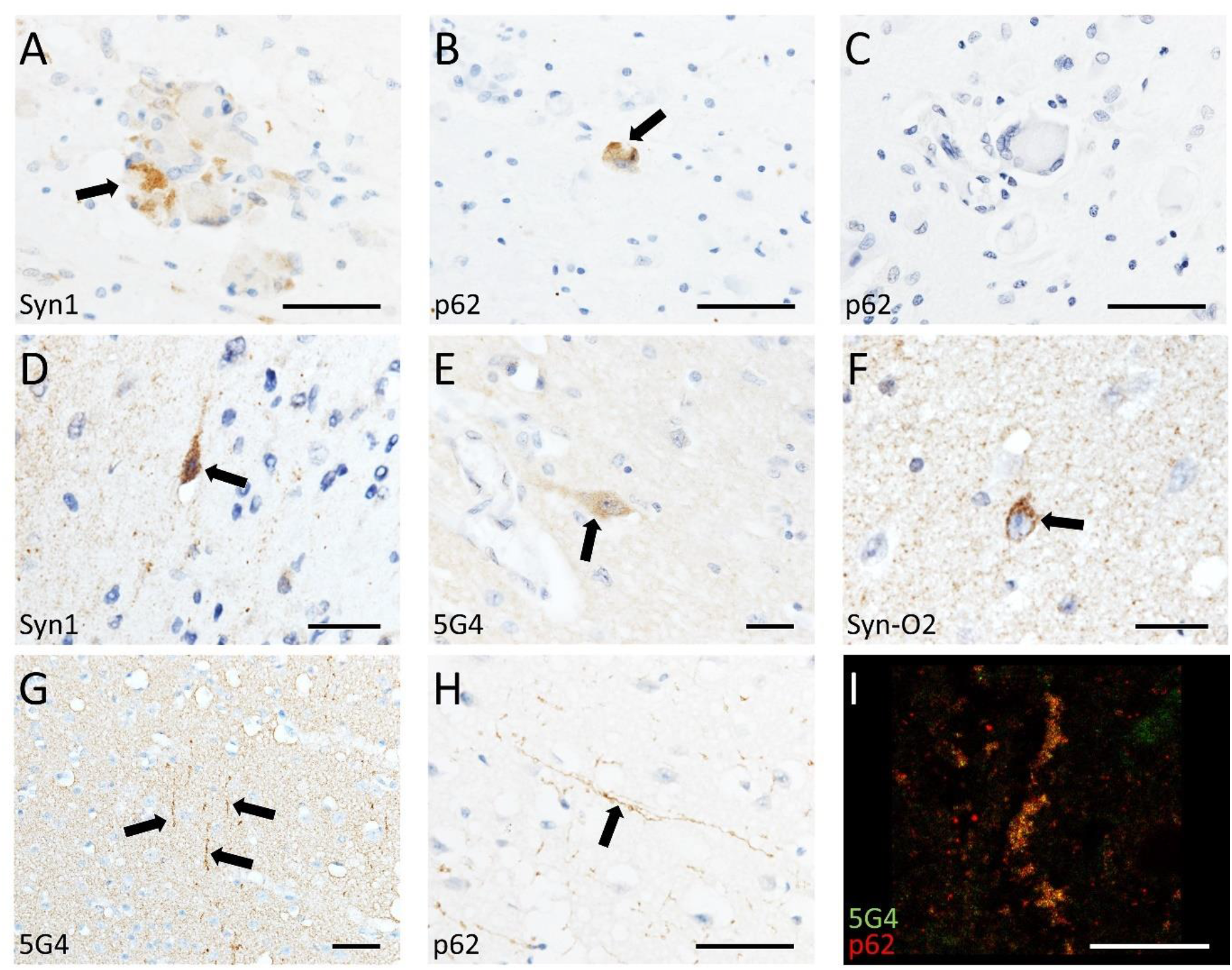
Photomicrographs of KD cases demonstrating α-synuclein-immunoreactive globoid cells (A) that were variably positive for the autophagic marker p62 (B-C). Occasional neurons manifested punctate α-synuclein staining (D-F) and neurites that were immunoreactive for α-synuclein and p62 were variably present throughout the cortex of all cases (G-I). 50 μm (A-C, G-H) 25 μm (D-F), 20 μm (I).

A sub-population of cortical pyramidal neurons in KD were labelled by the Syn1 antibody and, although α-synuclein can be observed as uniform perikaryal staining in some infant cortical neurons prior to the age of 12 months [9], neuronal α-synuclein in KD was present in both cases over 12 months of age and had a granular rather than uniform appearance (Figure 1D). A sub-set of KD pyramidal neurons were labelled by the LBD-associated conformation-specific α-synuclein antibodies Syn-O2 and 5G4 (Figure 1, E-F; Supplementary Figure 1 & 3). Additionally, long cortical neurites that were immunoreactive for p62/SQSTM1 and 5G4 were observed with varying frequency across cases (Figure 1, G-I; Supplementary Figure 1 & 4). We did not observe immunoreactivity for α-synuclein phosphorylated at serine 129 (pS129), which is associated with LBD, in any of the aforementioned structures.

The 5G4 and Syn-O2 antibodies can recognise early oligomeric forms of α-synuclein aggregates [10, 11] whilst pS129 is localised to the external surface of Lewy bodies [8], perhaps suggesting it is a late modification in α-synuclein aggregation. Therefore, as α-synuclein in KD was immunoreactive for 5G4 and Syn-O2 but not pS129, it is tempting to speculate that this indicates α-synuclein in KD is at an earlier stage of aggregation than that observed in mature Lewy bodies in LBD. Such a proposition is consistent with the rapid disease course of KD, ranging from 4-10 months in the present study, in contrast to the long prodromal phase of LBD of up to 20 years based on α-synuclein in colonic biopsies [12].

### Brain lysates from KD manifest high molecular weight α-synuclein species and are seed-competent on RT-QuIC

To further explore the nature of α-synuclein aggregates in KD we performed immunoblotting on two cases for which frozen tissue was available and compared these to four infant control cases and one LBD case as a synucleinopathy disease-control. We selected temporal cortex as it manifested α-synuclein pathology in both KD cases and evaluated grey and white matter separately. Whilst we did not observe consistent changes in total α-synuclein in KD or the LBD case, likely due to the abundance of monomeric species, we identified subtle increases in pS129 relative to total α-synuclein (Supplementary Figure 5). Dysregulated α-synuclein was also identified in both grey and white matter of KD cases, as shown by higher molecular weight bands of approximately 20 and 40 kDa that were remarkably similar in both KD cases (Figure 2, A-B). However, the higher molecular weight pS129 bands in KD cases differed from those in the LBD case included as a disease control, suggesting that although α-synuclein is dysregulated in KD, it differs from that observed in LBD (Figure 2, A-B and Supplementary Figure 5 & 6).

**Figure 2:**
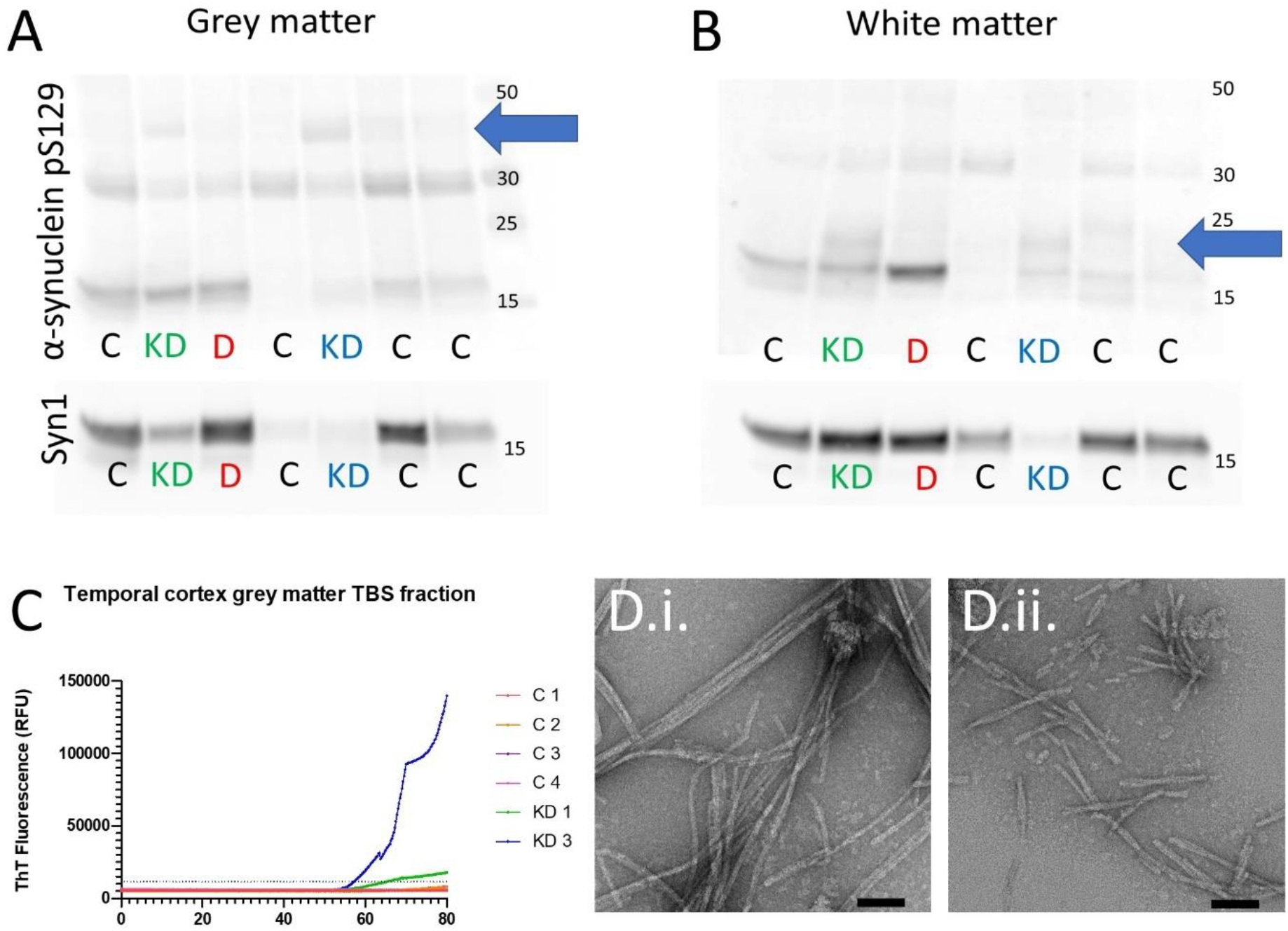
The TEAB-soluble fraction of KD temporal cortex grey matter lysates contained pS129 bands at approximately 40 kDa (blue arrow) that were not observed in control cases or the DLB case (A). In white matter samples, KD cases had additional bands of pS129 at approximately 20 kDa (blue arrow) that were not present in control or DLB cases (B). RT-QuIC of the TBS-soluble tissue fractions demonstrated that KD1 and KD3 grey matter gave positive reactions, defined as thioflavin-T fluorescence greater than five times the control standard deviation, at approximately 55 hours whilst no reaction was observed in white matter samples (C). Transmission electron microscopy of RT-QuIC end-products from grey matter of KD 1 (D.i.) and KD3 (D.ii.) confirmed the presence of fibrillar structures. It is notable that KD1 RT-QuIC end-products led to elongated fibrils of approximately 200-400 nm whilst KD3 end-products were typically 100-200 nm (E.i.-F.ii.). Scale bars = 50 nm (D.i. & D.ii.).

A key pathogenic quality ascribed to α-synuclein in LBD is its prion-like capacity, where a ‘seed’ of misfolded α-synuclein can induce α-synuclein monomers to misfold and aggregate into insoluble fibrillar forms [3]. As psychosine, the cytotoxic sphingolipid that accumulates in KD white matter, has been reported to induce α-synuclein misfolding and aggregation *in vitro* [6], we hypothesised that α-synuclein in KD would have prion-like qualities and would therefore be capable of seeding monomeric α-synuclein to misfold into fibrils. Therefore, we performed the real-time quaking-induced conversion (RT-QuIC) assay, where KD temporal cortex samples were mixed with recombinant α-synuclein and agitated, on the basis that if there is prion-like α-synuclein within the tissues they will seed the aggregation of the recombinant protein into fibrils detectable by the amyloid dye Thioflavin-T [13]. RT-QuIC identified that the aqueous fraction of the grey matter of both KD cases seeded aggregation into fibrils, indicating it has prion-like capacity (Figure 2, C), with similar intensity to LBD samples we have previously reported using the same assay [13]. In contrast, the detergent-soluble fraction of grey matter and both white matter tissue fractions did not give a positive RT-QuIC reaction (Supplementary Figure 7). Transmission electron microscopy of RT-QuIC end-products from the aqueous fraction of grey matter from both KD cases confirmed the presence of fibrils (Figure 2, D.i.-D.ii.). Our previous findings in LBD cases suggested both aqueous-soluble and detergent-soluble fractions had prion-like capacity and, as one would expect soluble conformers to precede the formation of detergent-soluble species, our observation of prion-like capacity limited to soluble α-synuclein in KD is consistent with the suggestion these aggregates are less mature than those observed in LBD.

Prion-like α-synuclein is a feature of an age-associated disease [13], thus its presence in infants with Krabbe disease likely indicates an association between altered sphingolipid metabolism and α-synuclein aggregation that may be relevant to the pathogenesis of LBD. A previous meta-analysis using over 19,000 PD cases reported variants in the *GALC* gene as a risk factor for PD [4] but we did not find an association in our smaller cohort of 2,414 DLB cases from previous studies [14, 15] (Supplementary Results 1-2), perhaps due to higher levels of concomitant Alzheimer-type pathology that influence the DLB clinical phenotype making it a less pure disorder of α-synuclein [16, 17]. However, we identified two heterozygous missense variants that were potentially deleterious (Combined Annotation Dependent Depletion score > 20) and only observed in DLB cases (Supplementary Results 3).

### Summary

For the first time, we report the presence of α-synuclein species with prion-like qualities in the brains of infants. These findings challenge the putative view that α-synuclein aggregation is merely an age-associated pathological alteration and instead suggest dysfunction of common biological pathways, such as sphingolipid metabolism, may underlie this phenomenon. Furthermore, the rapid disease course of KD indicates that the development of prion-like α-synuclein can occur in a more accelerated time-period than previously thought. Future studies are warranted to investigate the role of sphingolipid homeostasis in the context of LBD and the potential efficacy of anti-α-synuclein therapies for KD. Given the presently reported association between disrupted sphingolipid homeostasis and α-synuclein aggregation, future studies should evaluate whether LBD-associated changes in other sphingolipidoses with putative links to LBD, such as Tay-Sachs disease and Niemann-Pick Type C1, even in the absence of overt Lewy body pathology [10].

## Methods

### Port-mortem tissue preparation

*Post-mortem* tissue was prepared according to standardised brain bank protocols (Supplementary Methods 1). Ethical approval was granted by Newcastle Brain Tissue Resource Ethics Committee for KD1 and KD3 whilst approval for KD2 and KD4, which were obtained from the University of Maryland Brain Bank via the NIH NeuroBioBank (USA) and the Medical University of Vienna (Austria), was granted by Newcastle University Ethical Review Board (8130/2020 and 14201/2020, respectively). KD cases were included if they had clinical features consistent with infantile KD, alongside enzymatic or genetic evidence of β-galactocerebrosidase deficiency or *GALC* mutation, respectively, and typical neuropathological features of KD (globoid cells and demyelination). KD4 did not have genetic or enzymatic evidence of KD; however, they were included as they had typical clinical and pathological features of KD and a sibling exhibited reduced galactosylceramidase activity. Demographic information for all cases is included in Supplementary Table 1.

### Immunohistochemical and fluorescent staining

6 μm sections from frontal, temporal, parietal, and occipital cortex, in addition to samples from medulla, midbrain, and cerebellum, were obtained for neuropathological evaluation with haematoxylin and eosin (H&E), Syn1 and p62/SQSTM1 staining (Supplementary Table 2). KD4 only had tissue from frontal cortex, temporal cortex and medulla available. The evaluated regions were chosen based on their putative pathological involvement in LBD or KD. Additional α-synuclein antibodies (Syn-O2, 5G4 and pS129) were evaluated in cortex, based on observations from Syn1 staining (Supplementary Table 2). Antibody visualisation used the Menarini MenaPath X-Cell Linked Plus HRP detection kit, as per manufacturer’s instructions (Menarini Diagnostics, Berkshire, UK). Sections were evaluated on a Nikon Eclipse 90i microscope with DsFi1 camera coupled to a computer with NIS Elements software v3 (Nikon, Tokyo, Japan). Immunofluorescent staining was performed to identify co-localisation of proteins of interest and visualised with secondary antibodies conjugated to fluorophores (Supplementary Table 2). Fluorescence was evaluated on a Leica SP8 confocal microscope coupled to a computer with LASX software (Leica, Wetzlar, Germany).

### SDS-PAGE and western blot analysis

Temporal cortex (Brodmann area 20) was chosen for molecular analysis of α-synuclein as it contained sufficient tissue for downstream analyses and consistently contained globoid cells and neuritic and neuronal α-synuclein pathology in KD cases. Frozen tissue was only available for two KD cases (KD 1 and KD 3) and these were compared to four infant control cases and a DLB case which was included as a synucleinopathy disease control (Supplementary Table 1). Tissue was fractionated by centrifugation into aqueous- and detergent-soluble fractions (Supplementary Methods 2).

Protein titres were determined and normalised across samples using the Bradford assay (Thermo Scientific, Paisley, UK). For denaturing gel electrophoresis, samples were mixed with lithium dodecyl sulphate (LDS; NuPage, Thermo Scientific, Paisley, UK) and dithiothreitol (15 mM DTT, Sigma, St Louis, MO, USA) before boiling at 70°C for 10 minutes. Samples (10 μg/lane) were separated on 4-12% Bis-Tris gels in MOPS buffer and transferred to 0.45 μm nitrocellulose membranes via semi-dry conditions (iBlot 2, NuPage, Thermo Scientific, Paisley, UK). PageRuler Plus prestained protein ladders (3 μg/lane) were included as a known protein molecular weight standard.

α-synuclein was fixed to membranes by boiling for 10 minutes in PBS before membranes were washed in 0.05 % Tween 20 (Sigma) containing Tris-buffered saline (TBST, in mM; 50 Trizma base, 150 NaCl, pH = 7.6) and blocked for 1 h at RT in 5 % bovine serum albumin (BSA) containing TBST prior to overnight incubation at 4 °C with primary antibodies (Table 3) under agitation. Antibody binding was visualised with secondary antibodies (Table 3) conjugated to HRP and chemiluminescence using 1.25 mM luminol, 30 μM coumaric acid and 0.0015% hydrogen peroxide. Membranes were visualised on a Fuji LAS4000 with Fuji imaging software (Fuji LAS Image, Raytek, Sheffield, UK) and equal loading across lanes was confirmed via staining with Ponceau S (Sigma, St Louis, MO, USA).

### Real-time quaking-induced conversion (RT-QuIC)

Expression and purification of recombinant full-length α-synuclein was performed as previously, using the pRK172 plasmid containing cDNA for the human *SNCA* gene expressed in E. Coli BL21 (DE3) and purified using size exclusion and Mono Q anion exchange chromatography [16]. α-Synuclein was diluted in 20 mM Tris/HCl pH 7.4, 100 mM NaCl, and protein concentrations were determined using the bicinchoninic acid assay (BCA; Pierce, Waltham, MA, USA) and aliquots (300 μL of 1 mg/mL) were prepared and stored at −80 °C. Prior to use, the proteins were filtered (100 kDa spin filter) and the protein concentration was again determined by BCA assay.

Brain tissue lysates from KD1 and KD3 and from four infant control cases were homogenised in a rotator-stator homogeniser at 10% w/v on ice in TBS (20 mM Tris-HCl pH 7.4, 150 mM NaCl) and 5 mM EDTA with protease and phosphatase inhibitors (Thermo Fisher, Paisley, UK). Samples were centrifuged at 3000 × g at 4 °C for 30 min and the supernatant recovered as the TBS (aqueous)-soluble fraction. The pellet was then resuspended in CelLytic buffer (Sigma, St Louis, MO, USA), homogenized on ice, and centrifuged at 3000× g at 4 °C for 30 min with the supernatant recovered as the detergent-soluble fraction. The total protein concentration was measured in both fractions by BCA assay (Pierce, Thermo Fisher, Paisley, UK) and aliquots were prepared and stored at −80 °C.

The RT-QuIC assay was performed as previously [16]. The reaction buffer was composed of 100 mM piperazine-N,N’-bis(ethanesulfonic acid) (PIPES; pH 6.9), 0.1 mg/mL αSyn, and 10 μM ThT. Reactions were performed in triplicate in black 96-well plates with a clear bottom (Nunc, Thermo Fisher, Paisley, UK) with 85 μL of the reaction mix loaded into each well together with 15 μL of 0.1 mg/ml TBS-soluble or detergent-soluble fractions. The plate was then sealed with a sealing film (Thermo Fisher, Paisley, UK) and incubated in a BMG LABTECH FLUOstar OMEGA plate reader at 37 °C for 100 h with intermittent cycles of 1 min shaking (500 rpm, double orbital) and 15 min rest throughout the indicated incubation time. ThT fluorescence measurements, expressed as arbitrary relative fluorescence units (RFU), were taken with bottom reads every 15 min using 450 ± 10 nm (excitation) and 480 ± 10 nm (emission) wavelengths for 80 hours and a positive signal was defined as RFU more than 5 standard deviation units (RFU > 5 SD) above the mean of initial fluorescence. The sample was considered positive if all of the replicates were positive, otherwise the sample was classified as negative. RT-QuIC was performed twice to confirm positive results.

### Electron microscopy of RT-QuIC end-products

Samples of RT-QuIC end-products from KD cases were transferred in a 5 μL volume to a TEM grid (S162 Formvar/Carbon, 200 Mesh, agar scientific) to confirm the presence of fibrils. After 1 min, samples were fixed using 5 μL of 0.5% glutaraldehyde followed by one wash with 50 μL of ddH2O. Five microliters of 2% uranyl acetate were added to the grid, and after two minutes uranyl acetate was blotted off and grids were dried before placing sample holders before visualization. TEM images were taken from both KD cases to confirm the presence of fibrils.

## Acknowledgements

This manuscript is dedicated to the memory of all the infants whose tissues were used in this study and their families for their generosity and selflessness in agreeing to donate brain tissue for research.

This study was funded by an Alzheimer’s Research UK Northern Network pump priming grant and the Alzheimer’s Research UK Fellowship awarded to DE (ARUK-RF2018C-005). The work conducted by the El-Agnaf laboratory was supported by Qatar Biomedical Research Institute under the Internal Grant Program SF 2017-007. CH was funded by South Tees Hospitals NHS Foundation Trust. DK is funded by The Lewy Body Society (LBS/007/2020). TFO is supported by the Deutsche Forschungsgemeinschaft (DFG, German Research Foundation) under Germany’s Excellence Strategy - EXC 2067/1-390729940, and by SFB1286 (B8). WH is supported by the NIHR GOSH BRC. The views expressed are those of the author(s) and not necessarily those of the NHS, the NIHR or the Department of Health.

Tissue for controls and KD cases 1 and 3 was obtained from Newcastle Brain Tissue Resource, a UK Human Tissue Authority approved brain tissue repository funded in part by a grant from the UK Medical Research Council (G0400074), by NIHR Newcastle Biomedical Research Centre awarded to the Newcastle upon Tyne NHS Foundation Trust and Newcastle University, and by a grant from the Alzheimer’s Society and Alzheimer’s Research UK as part of the Brains for Dementia Research Project. Tissue for KD case 2 was obtained from the University of Maryland Brain Bank, USA, and KD case 4 from the Medical University of Vienna, Austria.

## References

1. Graziano, A.C. and V. Cardile, History, genetic, and recent advances on Krabbe disease. Gene, 2015. 555(1): p. 2–13.

2. Nicaise, A.M., E.R. Bongarzone, and S.J. Crocker, A microglial hypothesis of globoid cell leukodystrophy pathology. J Neurosci Res, 2016. 94(11): p. 1049–61.

3. Outeiro, T.F., et al., Dementia with Lewy bodies: an update and outlook. Mol Neurodegener, 2019. 14(1): p. 5.

4. Chang, D., et al., A meta-analysis of genome-wide association studies identifies 17 new Parkinson’s disease risk loci. Nat Genet, 2017. 49(10): p. 1511–1516.

5. Abdelkarim, H., et al., alpha-Synuclein interacts directly but reversibly with psychosine: implications for alpha-synucleinopathies. Sci Rep, 2018. 8(1): p. 12462.

6. Smith, B.R., et al., Neuronal inclusions of alpha-synuclein contribute to the pathogenesis of Krabbe disease. J Pathol, 2014. 232(5): p. 509–21.

7. Erskine, D., et al., Lipids, lysosomes and mitochondria: insights into Lewy body formation from rare monogenic disorders. Acta Neuropathol, 2021. 141(4): p. 511–526.

8. Shahmoradian, S.H., et al., Lewy pathology in Parkinson’s disease consists of crowded organelles and lipid membranes. Nat Neurosci, 2019. 22(7): p. 1099–1109.

9. Raghavan, R., et al., Alpha-synuclein expression in the developing human brain. Pediatr Dev Pathol, 2004. 7(5): p. 506–16.

10. Kumar, S.T., et al., How specific are the conformation-specific alpha-synuclein antibodies? Characterization and validation of 16 alpha-synuclein conformation-specific antibodies using well-characterized preparations of alpha-synuclein monomers, fibrils and oligomers with distinct structures and morphology. Neurobiol Dis, 2020. 146: p. 105086.

11. Vaikath, N.N., et al., Generation and characterization of novel conformation-specific monoclonal antibodies for alpha-synuclein pathology. Neurobiol Dis, 2015. 79: p. 81–99.

12. Stokholm, M.G., et al., Pathological alpha-synuclein in gastrointestinal tissues from prodromal Parkinson disease patients. Ann Neurol, 2016. 79(6): p. 940–9.

13. Poggiolini, I., et al., RT-QuIC Using C-Terminally Truncated α-Synuclein Forms Detects Differences in Seeding Propensity of Different Brain Regions from Synucleinopathies. Biomolecules, 2021. 11(6): p. 820.

14. Orme, T., et al., Analysis of neurodegenerative disease-causing genes in dementia with Lewy bodies. Acta Neuropathol Commun, 2020. 8(1): p. 5.

15. Guerreiro, R., et al., Investigating the genetic architecture of dementia with Lewy bodies: a two-stage genome-wide association study. Lancet Neurol, 2018. 17(1): p. 64–74.

16. Edison, P., et al., Amyloid load in Parkinson’s disease dementia and Lewy body dementia measured with [11C]PIB positron emission tomography. J Neurol Neurosurg Psychiatry, 2008. 79(12): p. 1331–8.

17. Walker, L., L. Stefanis, and J. Attems, Clinical and neuropathological differences between Parkinson’s disease, Parkinson’s disease dementia and dementia with Lewy bodies - current issues and future directions. J Neurochem, 2019. 150(5): p. 467–474.

