## Supplementary Data for "Prion-like α-synuclein pathology in the brains of infants: Krabbe disease as a novel seed-competent α-synucleinopathy"

*Tables*

Table 1: Demographic details of the cohort. “KD” signifies Krabbe disease, “SIDS” refers to sudden infant death syndrome, “N.D.” means not determined, “DLB” refers to dementia with Lewy bodies.

| Case ID | Sex | Diagnosis | Presentation | Disease duration | Age at death | Brain weight |
|---|---|---|---|---|---|---|
| KD 1 | Male | KD | Failure to thrive and developmental delay at 6 months, later developed frequent apnoea and vomiting, spastic quadriplegia, microcephaly and visual loss. Diagnosis by clinical impression and later confirmed by *post-mortem* demonstration of absent galactocerebrosidase activity in cortex. | 4 m | 10 m | 660g |
| KD 2 | Male | KD | Central hypotonia and head lag at 4 months, developmental delay and persistent crying noted at 5.5 months and later developed seizure-like activity. Diagnosis confirmed by reduced galactocerebrosidase activity. | 6 m | 10 m | N.D. |

| | | | | | | |
|---|---|---|---|---|---|---|
| KD 3 | Female | KD | Status epilepticus following chest infection at 2 months, followed by developmental delay and feeding difficulties. Later developed central hypotonia and spasticity, then epileptic encephalopathy. Diagnosis confirmed by reduced galactocerebrosidase activity and genetic testing indicated homozygous 30-kb deletion in exons 11 through 17 of *GALC* gene. | 10 m | 12 m | 690g |
| KD4 | Male | KD | Following vaccination at 5 months developed enteritis followed by restlessness and crying, followed by spastic tetraplegia, opisthotonos, apnoea and continuous crying. Later developed generalised myoclonus, aspiration episodes, bradycardia and pneumonia, against backdrop of progressive failure to thrive. Diagnosis was made by neuropathological examination but a sibling later demonstrated reduced galactocerebrosidase activity. | 10 m | 15 m | 730g |
| C 1 | Male | SIDS | Parent awoke to find infant unresponsive in their arms, pronounced dead at arrival to hospital. SIDS diagnosed due to absence of obvious cause of death on *post-mortem* examination. | N.A. | 3 m | 640g |

| | | | | | | |
|---|---|---|---|---|---|---|
| C 2 | Male | Hypoxic-ischaemic encephalopathy | Hypoxia-ischaemia due to prolonged labour, died after one month in the Special Care Baby Unit. | 1 m | 1 m | N.D. |
| C 3 | Female | SIDS | Parents discovered deceased in bed. SIDS diagnosed due to absence of obvious cause of death on *post-mortem* examination. | N.A. | 20 m | N.D. |
| C 4 | Male | SIDS | Parents discovered deceased in cot. SIDS diagnosed due to absence of obvious cause of death on *post-mortem* examination. | N.A. | 2 m | 570g |
| DLB | Male | DLB | Presented with visual hallucinations and cognitive fluctuations, diagnosed with dementia with Lewy bodies. | 4 y | 81 y | 1331g |

Table 2: Antibodies used for immunohistochemistry and immunofluorescence

| IHC antibody | Isotype | Manufacturer | Dilution | Antigen retrieval |
|---|---|---|---|---|
| Anti-α-synuclein Syn1/Clone 42 | Mouse IgG1 | BD Biosciences #610786 | 250 µg/ml | Citrate pH 6, formic acid |
| Anti-oligomeric α-synuclein (Syn-O2) | Mouse IgG1 | El-Agnaf laboratory | 100 ng/ml | Citrate pH 6, formic acid |
| Anti-aggregated α-synuclein (5G4) | Mouse IgG1 | Merck MABN389 | 200 ng/ml | Citrate pH 6, formic acid |
| Anti-α-synuclein pS129 | Rabbit IgG | Abcam ab51253 | 6 µg/ml | Citrate pH 6, formic acid |
| Anti-SQSTM1/p62 | Mouse IgG1 | BD Biosciences #610832 | 8 µg/ml | Citrate pH 6 |

| IF antibody | Isotype | Manufacturer | Dilution | Secondary antibody |
|---|---|---|---|---|
| Anti-α-synuclein Syn1/Clone 42 | Mouse IgG1 | BD Biosciences #610786 | 1 µg/ml | Goat anti-mouse IgG1-AF488 (Thermo #A-21121) |
| Anti-oligomeric α-synuclein (Syn-O2) | Mouse IgG1 | El-Agnaf laboratory | 500 ng/ml | Goat anti-mouse IgG1-AF488 (Thermo #A-21121) |

| | | | | |
|---|---|---|---|---|
| Anti-aggregated α-synuclein (5G4) | Mouse IgG1 | Merck MABN389 | 200 ng/ml | Goat anti-mouse IgG1-AF488 (Thermo #A-21121) |
| Anti-SQSTM1/p62 | Rabbit IgG | Abcam ab207305 | 2 μg/ml | Goat anti-rabbit IgG-AF546 (Thermo #A-11035) |
| Anti-NeuN | Mouse IgG2b | Abcam ab104224 | 5 μg/ml | Goat anti-mouse IgG2b-647 (Thermo #A-21242) |

Table 3: Antibodies used for western blot analysis

| Antibody | Manufacturer | Dilution | Secondary antibody | Dilution |
|---|---|---|---|---|
| Anti-α-synuclein Syn1/Clone 42 | BD Biosciences #610786 | 250 ng/ml | Goat anti-mouse | 1:5,000 |
| Anti-α-synuclein pS129 | Abcam ab51253 | 3 μg/ml | Goat anti-rabbit | 1:2,500 |

*Figures*

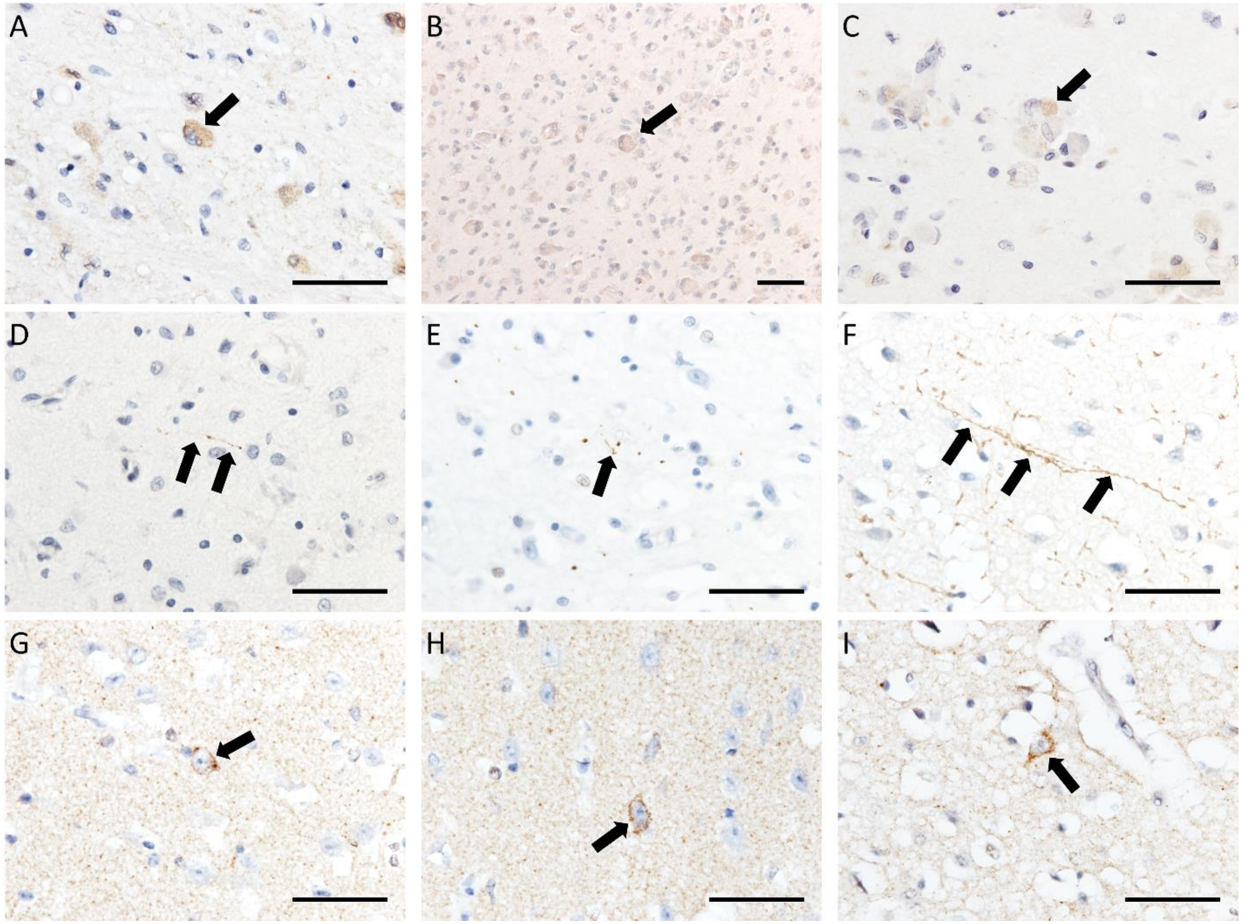


Figure 1: Photomicrographs from KD cases demonstrating Syn1 immunoreactivity in globoid cells (A-C), p62+ neurites (D-F), and Syn-O2 in pyramidal neurons (G-I). Cases are KD1 (E & I), KD2 (B), KD3 (C, F & G). KD4 (A, D & H), scale bars are 50 µm.

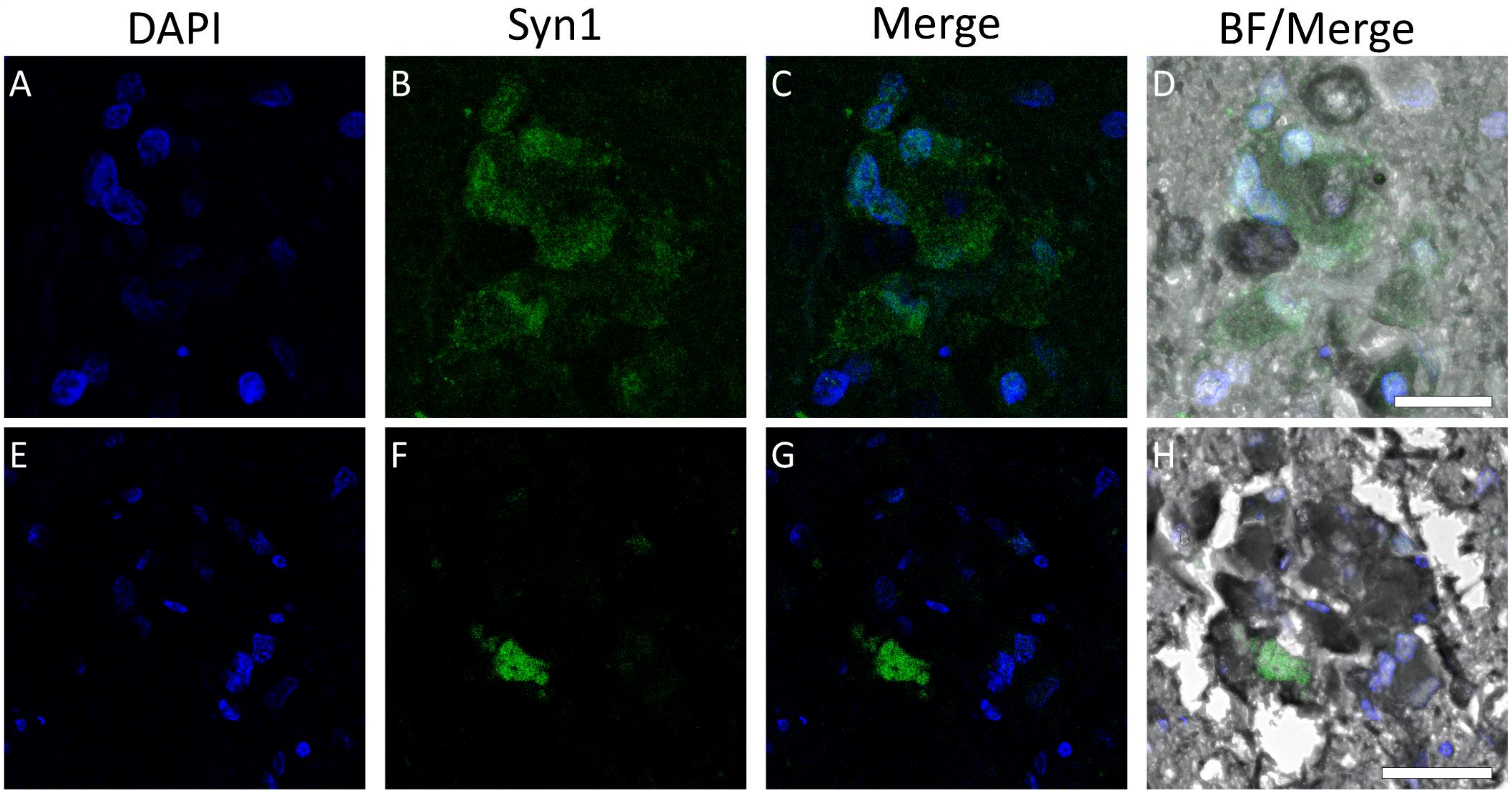


Figure 2: Immunofluorescent images from KD cases demonstrating Syn1 localised with globoid cell clusters in the medial lemniscus of KD1 following Sudan Black B treatment, confirming that α-synuclein is not non-specific recognition of lipid deposits. Scale bars = 40 μm.

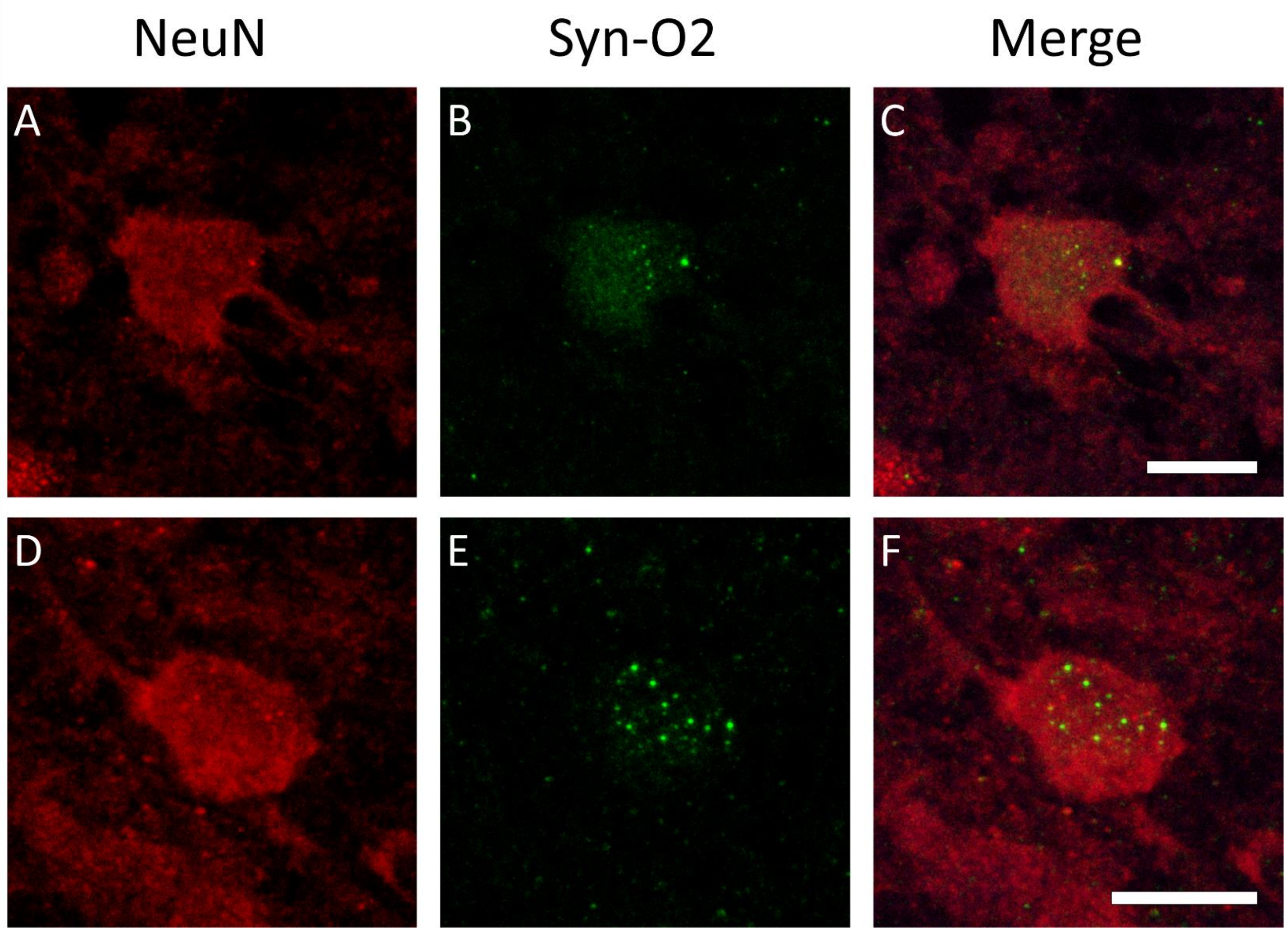


Figure 3: Immunofluorescent images demonstrating Syn-O2 punctae within temporal cortex cell bodies in KD1 (A-C) and KD3 (D-F). Scale bars = 10 μm.

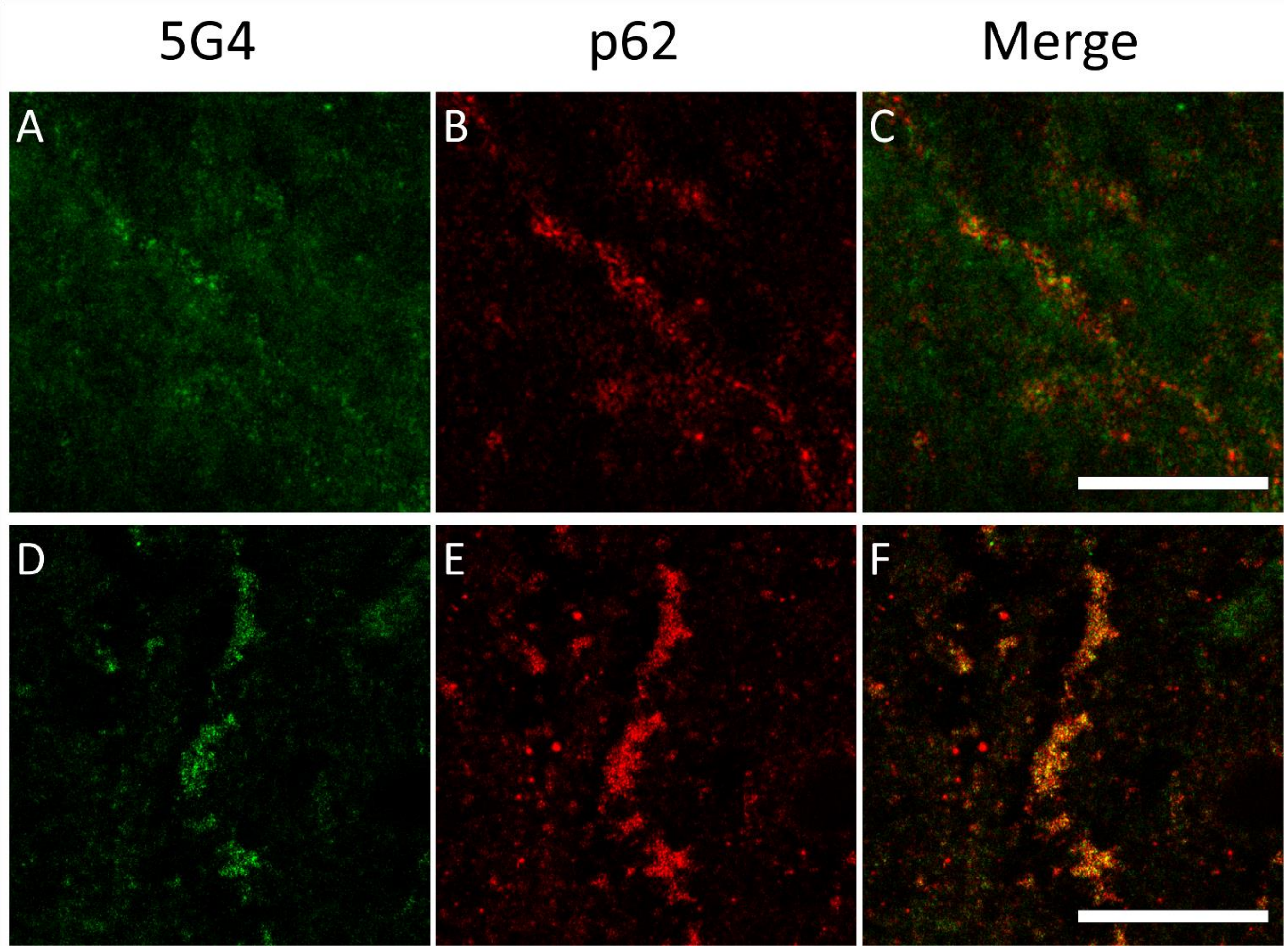


Figure 4: Fluorescent images demonstrating 5G4-immunoreactivity in p62+ neurites in KD3 (A-C) and KD2 (D-F). Scale bars = 10 μm (A-C) and 20 μm (D-F).

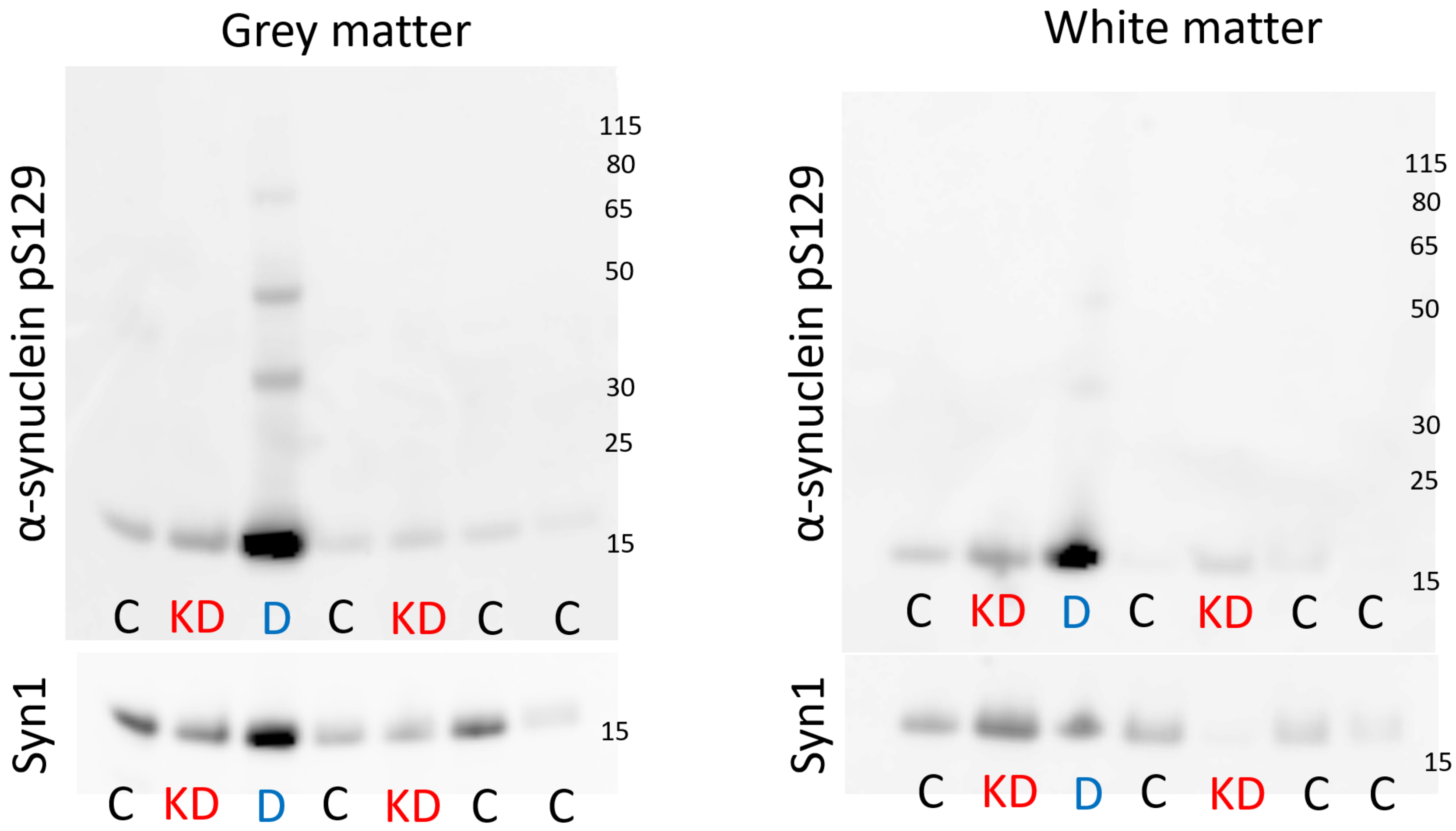


Figure 5: Western blot of crude temporal cortex lysates immunostained with pS129 (top) and Syn1 (bottom), KD cases (red) are indistinguishable from controls (black). In contrast, the DLB case (blue) demonstrates higher levels of pS129 in both grey and white matter, though high molecular weight bands are only obvious in grey matter. In all cases, Syn1 is variable across cases and particularly marked in DLB grey matter.

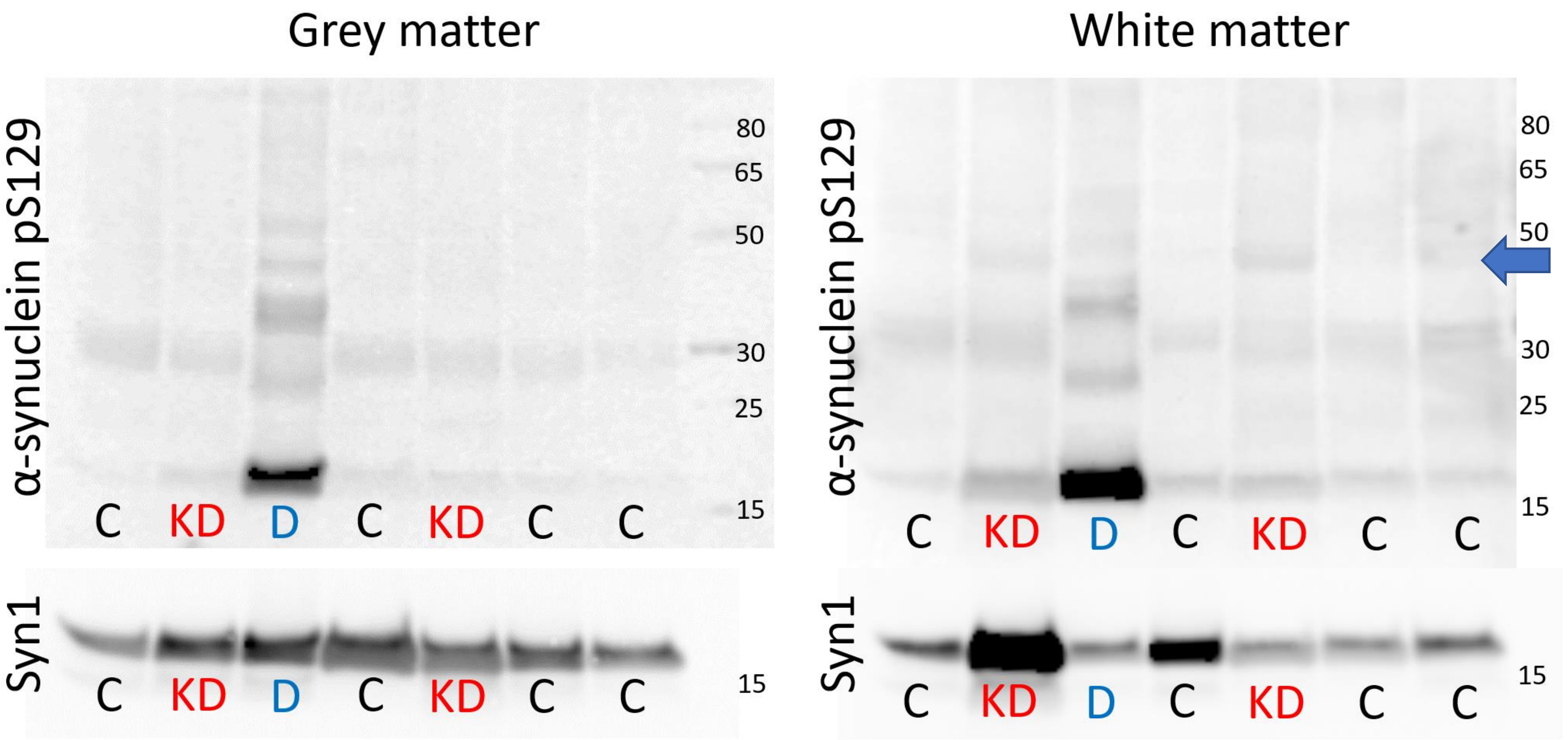


Figure 6: Western blot of Triton X-100-soluble tissue lysates immunostained with pS129 (top) and Syn1 (bottom). KD cases (red) have higher molecular weight bands of pS129 at approximately 45 kDa in white matter (blue arrow) but are no different from controls in grey matter. The DLB case (blue) has numerous high molecular weight pS129 bands in both grey and white matter. Syn1 is not different in DLB compared to control in grey or white matter but KD2 has strikingly high levels of Syn1 in white matter. It is notable that KD2 had very high densities of Syn1-immunoreactive globoid cells in this area.

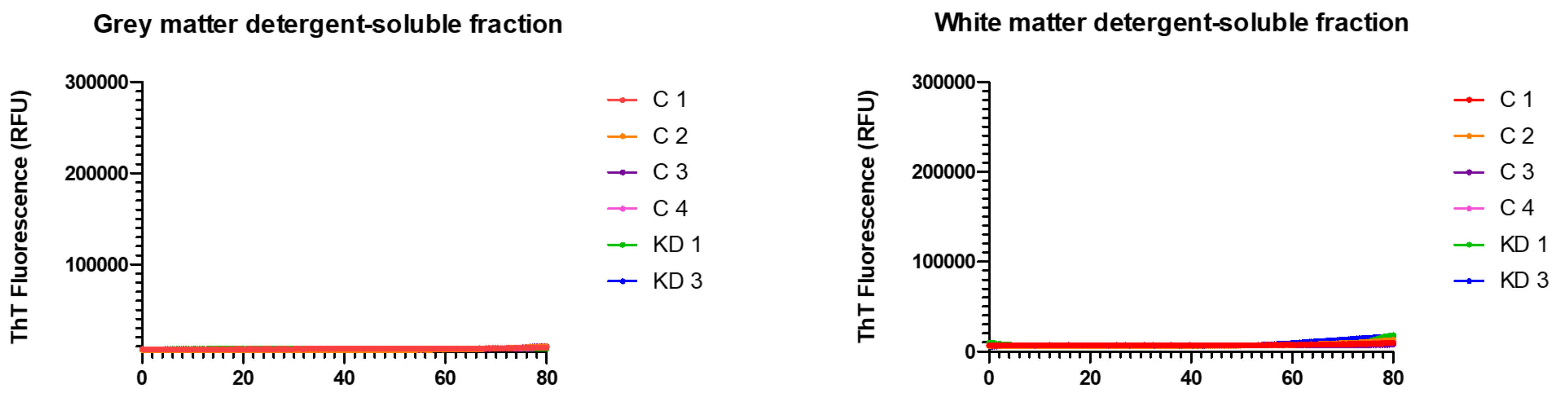


Figure 7: RT-QuIC data demonstrating that the detergent soluble fractions of both grey and white matter did not demonstrate a positive reaction with the α-synuclein RT-QuIC assay.

*Methods*

Supplementary Methods 1: Post-mortem tissue preparation

At autopsy, the brainstem was removed at the level of the red nucleus and the cerebrum hemisected through the corpus callosum. The right hemisphere was immersion-fixed in neutral-buffered formalin whilst the left hemisphere was sliced into approximately 1 cm coronal slices and snap frozen at -120°C prior to storage at -80°C. After five weeks fixation, the right hemisphere was sub-dissected for neuropathological evaluation and processed for paraffin wax embedding.

Supplementary Methods 2: Fractionation of tissue for immunoblotting

Approximately 400 mg of grey matter from the inferior temporal gyrus, in addition to 400 mg of underlying white matter, was dissected from tissue slices at -20°C. Small samples were retained for real-time quaking-induced conversion (RT-QuIC) analysis, whilst the remainder was suspended in 0.2 M TEAB buffer supplemented with Complete protease inhibitors and PhoStop tablets (Roche, Basel, Switzerland) and homogenised for 30 seconds in a rotator-stator homogeniser. Approximately 200 µl was removed from each sample as the crude tissue lysate.

Samples were centrifuged at 16,000 rcf for 30 minutes at 4°C with the supernatant recovered as the soluble tissue fraction. Samples were then washed by resuspension in 1:10 w/v Tris-HCl buffer (50 mM Tris-HCl, 150 mM sodium chloride, 2 mM EDTA) and centrifuged twice at 16,000 rcf for 30 minutes at 4°C. The pellet was resuspended 1:5 w/v in Tris-HCl buffer containing 5% SDS, vortexed, and incubated at room temperature for 5 minutes before centrifugation at 16,000 rcf for 30 minutes at 10°C, with the supernatant recovered as the SDS-soluble fraction.

Supplementary Methods 3: *GALC* gene and genetic risk of DLB

This analysis utilized data described in two previous studies: an exome sequenced cohort of 1,118 DLB patients [1] and a cohort of whole-genome genotyped samples described in [2] (Table 4). The data described here has been lifted over to the most recent genome build (GRCh38/hg38). The genotyping data were imputed on the TOPMed platform [3].

Table 4: Cohort descriptions. Whole exome sequencing (WES) and whole genome genotyped (WGG) data.

| Cohort Name | Publication | Data format | Cases | Controls | Average Age |
|---|---|---|---|---|---|
| DLB exomes | [1] | WES | 1,118 | 0 | 78.8 |
| DLB WGG | [2] | WGG | 1,296 | 4,379 | 79.64 |

### Quality control

Individual datasets were subject to variant QC conducted primarily in bcftools version 1.12 [4]. All datasets were aligned to the latest reference genome panel (GRCh38/hg38). Variant annotations were generated using SNPeff [5]. Variant and sample QC involved removing samples with low variant quality (DP < 8, GQ < 20, and missingness > 0.15) and discordant allelic depth and genotypes.

### Single variant association

The association to disease status of both rare and common variants was assessed using logistic and Firth regression. Logistic regression was performed on common (MAF > 0.05) variants using PLINK 2.0 [6] using the first 10 principal components (PCs) as covariates. Rare variant (MAF < 0.05) association was conducted using Firth regression, given its superior ability to control for type I errors [7, 8]. This test was performed using the implementation available in rvtests [9], also using the first 10 PCs as covariates.

Since no controls were available to complement the WES data, Gnomad allele counts from the non-Finnish European population, specifically those taken from the non-neurologically affected subset (https://gnomad.broadinstitute.org/about), were used in a fisher exact test to determine significance of variants. This test was implemented in R using the fisher.test() function.

### Gene burden analysis

Gene burden tests were conducted using SkatO as available through rvtests [9]. The first 10 PCs were used as covariates. The burdens of missense, high impact, and missense plus loss of function variants were conducted separately. Variant functional impact was determined using SNPeff [5].

*Results*

Supplementary Results 1: Single variant associations

No significant association was detected between common variants within the *GALC* gene region and disease status (Fig 8). We also detected no association between rare variants within the *GALC* gene region (Figure 9). The most significant result was a variant, rs561184126 (14-87988525-T-TA), from the fisher test in the exomes with a p-value of 0.003 and an OR of 53.83.

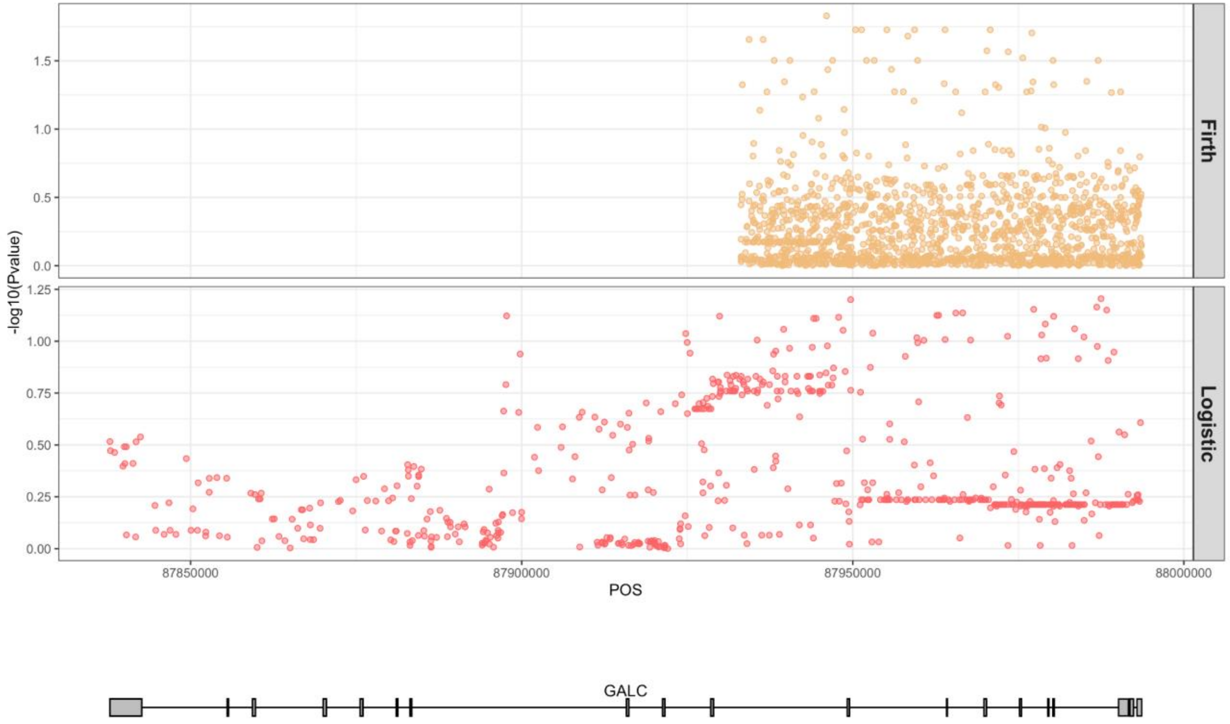


Figure 8. Logistic (bottom) and Firth (top) regression results for the whole genome

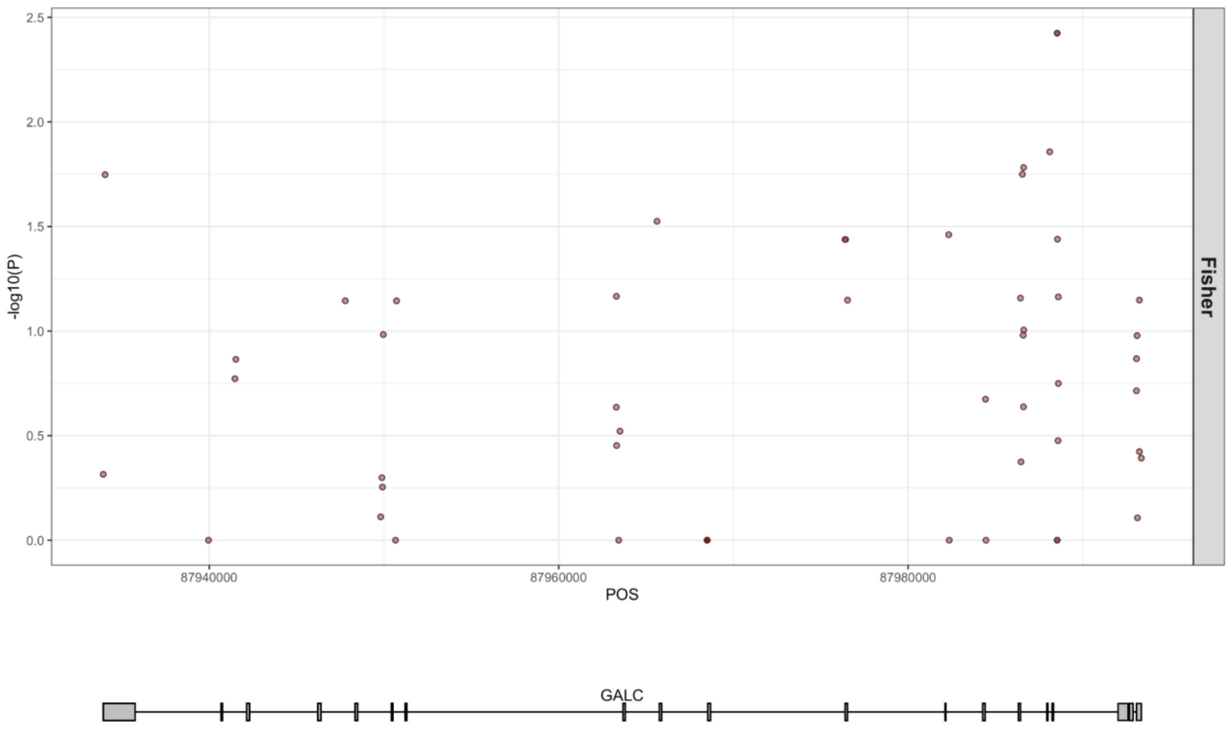


Figure 9. Fisher exact test results comparing allele counts in DLB exomes to those in non-neurologically affected non-Finnish Europeans from the Gnomad database

Supplementary Results 2: Gene burden analysis

Gene burden analysis demonstrated no significant associations in SkatO (Table 5).

Table 5: SkatO gene burden results for exome and genotyping data

| | Variant | NumVar | NumPolyVar | Q | rho | pvalue |
|---|---|---|---|---|---|---|
| DLB Exomes vs ADSP Exome Controls | High Impact | 14 | 3 | NA | NA | NA |
| | Missense + LOF | 112 | 33 | 198372 | 0 | 0.186102 |
| | Missense | 98 | 30 | 110412 | 1 | 0.062663 |
| DLB WGG | High Impact | 3 | 2 | 22.0663 | 1 | 0.668939 |
| | Missense + LOF | 48 | 33 | 6242.71 | 1 | 0.474872 |
| | Missense | 45 | 31 | 16177.9 | 1 | 0.144063 |

Supplementary Results 3: Prioritised rare variants

No high-impact variants were identified in the genotyping data. Two heterozygous missense variants were found only in DLB cases (1 case each). The first of these variants, rs777305549, had a CADD score of 22.5 and a frequency of 1.55E-05 in the Gnomad Non-Finnish European population. The second of these variants, rs371523347, was reported on ClinVar with uncertain significance for galactosylceramide beta-galactosidase deficiency. This variant had a frequency of 9.30E-05 in the NFE Gnomad population, and a CADD score of 32.

The exome data similarly did not contain any case-specific homozygous variants. The most notable variants from this dataset were four missense, heterozygous variants with CADD scores > 20. None of these variants were described on ClinVar (Table 6).

Table 6: The most interesting prioritized variants from exome and genotyping data. Positions given in hg38.

| **SNP ID** | **Position** | **REF/ALT** | **Amino Acid Change** | **Gnomad MAF NFE** | **CADD** | **ClinVarReport** | **Data source** |
|---|---|---|---|---|---|---|---|
| rs777305549 | 87950702 | T/A | p.Asn403Ile | 1.55E-05 | 22.5 | NA | WGG |
| rs371523347 | 87988513 | C/T | p.Arg69Gln | 9.30E-05 | 32 | uncertain significance for galactosylceramide beta-galactosidase deficiency | WGG |
| chr14:87984483:A:G | 87984483 | A/G | p.Tyr165His | NA | 31 | NA | WES |
| rs772928724 | 87993063 | A/C | p.Cys34Trp | 7.75E-05 | 27.6 | NA | WES |

| | | | | | | | |
|---|---|---|---|---|---|---|---|
| chr14:87993083:C:G | 87993083 | C/G | p.Ala28Pro | NA | 22.6 | NA | WES |
| rs372285275 | 87993100 | C/T | p.Gly22Asp | 1.55E-05 | 22.8 | NA | WES |

### References


1. Orme, T., et al., *Analysis of neurodegenerative disease-causing genes in dementia with Lewy bodies.* Acta Neuropathol Commun, 2020. **8**(1): p. 5.
2. Guerreiro, R., et al., *Investigating the genetic architecture of dementia with Lewy bodies: a two-stage genome-wide association study.* Lancet Neurol, 2018. **17**(1): p. 64-74.
3. Taliun, D., et al., *Sequencing of 53,831 diverse genomes from the NHLBI TOPMed Program.* Nature, 2021. **590**(7845): p. 290-299.
4. Danecek, P., et al., *Twelve years of SAMtools and BCFtools.* Gigascience, 2021. **10**(2).
5. Cingolani, P., et al., *A program for annotating and predicting the effects of single nucleotide polymorphisms, SnpEff: SNPs in the genome of Drosophila melanogaster strain w1118; iso-2; iso-3.* Fly (Austin), 2012. **6**(2): p. 80-92.
6. Chang, C.C., et al., *Second-generation PLINK: rising to the challenge of larger and richer datasets.* Gigascience, 2015. **4**: p. 7.
7. Ma, C., et al., *Recommended joint and meta-analysis strategies for case-control association testing of single low-count variants.* Genet Epidemiol, 2013. **37**(6): p. 539-50.
8. Wang, X., *Firth logistic regression for rare variant association tests.* Front Genet, 2014. **5**: p. 187.
9. Zhan, X., et al., *RVTESTS: an efficient and comprehensive tool for rare variant association analysis using sequence data.* Bioinformatics, 2016. **32**(9): p. 1423-6.

*Supplementary author list*

*International DLB Genetics Consortium Author list:*

**USA:** Jose Bras (Department of Neurodegenerative Science, Van Andel Institute, Grand Rapids), Rita Guerreiro (Department of Neurodegenerative Science, Van Andel Institute, Grand Rapids), Celia Kun-Rodrigues (Department of Neurodegenerative Science, Van Andel Institute, Grand Rapids), Andrew Singleton (Laboratory of Neurogenetics, National Institutes on Aging, NIH, Bethesda, MD, USA), Dena Hernandez (Laboratory of Neurogenetics, National Institutes on Aging, NIH, Bethesda, MD, USA), Owen A. Ross (Department of Neuroscience, Mayo Clinic, Jacksonville, FL, USA), Dennis W. Dickson (Department of Neuroscience, Mayo Clinic, Jacksonville, FL, USA), Neill Graff-Radford (Department of Neurology, Mayo Clinic, Jacksonville, FL, USA), Tanis J. Ferman (Department of Psychiatry and Department of Psychology, Mayo Clinic, Jacksonville, FL, USA), Ronald C. Petersen (Neurology Department, Mayo Clinic, Rochester, MN, USA), Brad F. Boeve (Neurology Department, Mayo Clinic, Rochester, MN, USA), Michael G. Heckman (Division of Biomedical Statistics and Informatics, Mayo Clinic, Jacksonville, FL, USA), John Q. Trojanowski (Department of Pathology and Laboratory Medicine, Center for Neurodegenerative Disease Research, Perelman School of Medicine at the University of Pennsylvania, 3600 Spruce Street, Philadelphia, USA), Vivianna Van Deerlin (Department of Pathology and Laboratory Medicine, Center for Neurodegenerative Disease Research, Perelman School of Medicine at the University of Pennsylvania, 3600 Spruce Street, Philadelphia, USA), Nigel J. Cairns (Knight Alzheimers Disease Research Center, Department of Neurology, Washington University School of Medicine, Saint Louis, MO, USA), John C. Morris (Knight Alzheimers Disease Research Center, Department of Neurology, Washington University School of Medicine, Saint Louis, MO, USA), David J. Stone (Genetics and Pharmacogenomics, Merck and Co, West Point, Pennsylvania, USA), John D. Eicher (Genetics and Pharmacogenomics, Merck and Co, Boston, MA, USA), Lorraine Clark (Taub Institute for Alzheimer Disease and the Aging Brain and Department of Pathology and Cell Biology, Columbia University, New York, NY, USA), Lawrence S Honig (Taub Institute for Alzheimer Disease and the Aging Brain and Department of Pathology and Cell Biology, Columbia University, New York, NY, USA),

Karen Marder (Taub Institute for Alzheimer Disease and the Aging Brain and Department of Pathology and Cell Biology, Columbia University, New York, NY, USA), Geidy E. Serrano (Banner Sun Health Research Institute, 10515 W Santa Fe Drive, Sun City, AZ 85351, USA), Thomas G. Beach (Banner Sun Health Research Institute, 10515 W Santa Fe Drive, Sun City, AZ 85351, USA), Douglas Galasko (Department of Neurosciences, University of California, San Diego, La Jolla, CA, United States; Veterans Affairs San Diego Healthcare System, La Jolla, CA, United States), Eliezer Masliah (Division of Neurosciences and Laboratory of Neurogenetics, National Institute on Aging/NIH, Bethesda MD 20892-9205, USA).

**UK:** John Hardy (UK Dementia Research Institute (UK DRI) at UCL, London, UK; Department of Neurodegenerative Disease, UCL Institute of Neurology, London, UK), Lee Darwent (UK Dementia Research Institute (UK DRI) at UCL, London, UK; Department of Neurodegenerative Disease, UCL Institute of Neurology, London, UK), Olaf Ansorge (Nuffield Department of Clinical Neurosciences, Oxford Parkinson's Disease Centre, University of Oxford, Oxford, UK), Laura Parkkinen (Nuffield Department of Clinical Neurosciences, Oxford Parkinsons Disease Centre, University of Oxford, Oxford, UK), Kevin Morgan (Human Genetics, School of Life Sciences, University of Nottingham, Nottingham, UK), Kristelle Brown (Human Genetics, School of Life Sciences, University of Nottingham, Nottingham, UK), Anne Braae (Human Genetics, School of Life Sciences, University of Nottingham, Nottingham, UK), Imelda Barber (Human Genetics, School of Life Sciences, University of Nottingham, Nottingham, UK), Claire Troakes (Department of Basic and Clinical Neuroscience and Institute of Psychiatry, Psychology and Neuroscience, Kings College London, London, UK), Safa Al-Sarraj (Department of Basic and Clinical Neuroscience and Institute of Psychiatry, Psychology and Neuroscience, Kings College London, London, UK), Tom Warner (Queen Square Brain Bank, Department of Molecular Neuroscience, UCL Institute of Neurology, London, UK), Tammaryn Lashley (Queen Square Brain Bank, Department of Molecular Neuroscience, UCL Institute of Neurology, London, UK), Janice Holton (Queen Square Brain Bank, Department of Molecular Neuroscience, UCL Institute of Neurology, London, UK), Yaroslau Compta (Queen Square Brain Bank, Department of Molecular

Neuroscience, UCL Institute of Neurology, London, UK), Tamas Revesz (Queen Square Brain Bank, Department of Molecular Neuroscience, UCL Institute of Neurology, London, UK), Andrew Lees (Queen Square Brain Bank, Department of Molecular Neuroscience, UCL Institute of Neurology, London, UK), Henrik Zetterberg (UK Dementia Research Institute (UK DRI) at UCL, London, UK; Department of Neurodegenerative Disease, UCL Institute of Neurology, London), Valentina Escott-Price (MRC Centre for Neuropsychiatric Genetics and Genomics, School of Medicine, Cardiff University, Cardiff, UK), Stuart Pickering-Brown (Institute of Brain, Behaviour and Mental Health, Faculty of Medical and Human Sciences, University of Manchester, Manchester, UK), David Mann (Institute of Brain, Behaviour and Mental Health, Faculty of Medical and Human Sciences, University of Manchester, Manchester, UK), Peter St. George-Hyslop (Department of Clinical Neurosciences, Cambridge Institute for Medical Research, University of Cambridge, Cambridge, UK).

**Canada:** Ekaterina Rogaeva (Tanz Centre for Research in Neurodegenerative Diseases and department of Medicine, University of Toronto, Ontario, Canada), Peter St. George-Hyslop (Tanz Centre for Research in Neurodegenerative Diseases and department of Medicine, University of Toronto, Ontario, Canada).

**Spain:** Jordi Clarimon (Memory Unit, Department of Neurology, IIB Sant Pau, Hospital de la Santa Creu i Sant Pau, Universitat Autonoma de Barcelona, Barcelona, Spain; Centro de Investigacion Biomedica en Red en Enfermedades Neurodegenerativas (CIBERNED), Instituto de Salud Carlos III, Madrid, Spain), Alberto Lleo (Memory Unit, Department of Neurology, IIB Sant Pau, Hospital de la Santa Creu i Sant Pau, Universitat Autonoma de Barcelona, Barcelona, Spain; Centro de Investigacion Biomedica en Red en Enfermedades Neurodegenerativas (CIBERNED), Instituto de Salud Carlos III, Madrid, Spain), Estrella Morenas-Rodriguez (Memory Unit, Department of Neurology, IIB Sant Pau, Hospital de la Santa Creu i Sant Pau, Universitat Autonoma de Barcelona, Barcelona, Spain; Centro de Investigacion Biomedica en Red en Enfermedades Neurodegenerativas (CIBERNED), Instituto de Salud Carlos III, Madrid, Spain), Pau Pastor (Memory Unit, Department of Neurology, University Hospital Mutua de Terrassa, University of Barcelona, and Fundacio de Docencia I Recerca Mutua de Terrassa, Terrassa, Barcelona, Spain. Centro de

Investigacion Biomedica en Red Enfermedades Neurdegenerativas (CIBERNED), Madrid, Spain), Monica Diez-Fairen (Memory Unit, Department of Neurology, University Hospital Mutua de Terrassa, University of Barcelona, and Fundacio de Docencia I Recerca Mutua de Terrassa, Terrassa, Barcelona, Spain. Centro de Investigacion Biomedica en Red Enfermedades Neurdegenerativas (CIBERNED), Madrid, Spain), Miquel Aquilar (Memory Unit, Department of Neurology, University Hospital Mutua de Terrassa, University of Barcelona, and Fundacio de Docencia I Recerca Mutua de Terrassa, Terrassa, Barcelona, Spain. Centro de Investigacion Biomedica en Red Enfermedades Neurdegenerativas (CIBERNED), Madrid, Spain), Yaroslau Compta (Movement Disorders Unit, Neurology Service, Clinical Neuroscience Institute (ICN), Hospital Clinic, University of Barcelona, IDIBAPS, Barcelona, Spain).

**Australia:** Claire Shepherd (Neuroscience Research Australia, Sydney, Australia and School of Medical Sciences, Faculty of Medicine, University of New South Wales, Sydney, Australia), Glenda M. Halliday (Brain and Mind Centre, Sydney Medical School, The University of Sydney, Sydney, Australia and Neuroscience Research Australia, Sydney, Australia and School of Medical Sciences, Faculty of Medicine, University of New South Wales,Sydney, Australia).

**Finland:** Pentti J. Tienari (Molecular Neurology, Research Programs Unit, University of Helsinki, Department of Neurology, Helsinki University Hospital, Helsinki, Finland), Liisa Myllykangas (Department of Pathology, University of Helsinki and Helsinki University Hospital, Helsinki, Finland), Minna Oinas (Department of Neuropathology and Neurosurgery, Helsinki University Hospital and University of Helsinki, Helsinki, Finland).

**Portugal:** Isabel Santana (Neurology Service, University of Coimbra Hospital, Coimbra, Portugal).

**France:** Suzanne Lesage (Inserm U1127, CNRS UMR7225, Sorbonne Universites, UPMC Univ Paris 06, UMR and S1127, Institut du Cerveau et de la Moelle epiniere, Paris, France).

**Sweden:** Henrik Zetterberg (Clinical Neurochemistry Laboratory, Institute of Neuroscience and Physiology, Sahlgrenska Academy at the University of Gothenburg, Molndal, Sweden), Elisabet Londos (Clinical Memory Research Unit, Institution of Clinical Sciences Malmo, Lund University, Sweden).

**The Netherlands:** Afina Lemstra (Department of Neurology and Alzheimer Center, Neuroscience Campus Amsterdam, VU University Medical Center, Amsterdam, the Netherlands).

### Acknowledgements

For the neuropathologically confirmed samples from Australia, tissues were received from the Sydney Brain Bank which is supported by Neuroscience Research Australia and the University of New South Wales. We would like to thank the South West Dementia Brain Bank (SWDBB) for providing brain tissue for this study. The SWDBB is supported by BRACE (Bristol Research into Alzheimer's and Care of the Elderly), Brains for Dementia Research and the Medical Research Council. The brain samples and/or bio samples were obtained from The Netherlands Brain Bank, Netherlands Institute for Neuroscience, Amsterdam (open access: www.brainbank.nl). All Material has been collected from donors for or from whom a written informed consent for a brain autopsy and the use of the material and clinical information for research purposes had been obtained by the NBB. This study was also partially funded by the Wellcome Trust, Medical Research Council and Canadian Institutes of Health Research (Dr. St. George-Hyslop). Work from Dr. Compta was supported by the CERCA Programme / Generalitat de Catalunya, Barcelona, Catalonia, Spain. This study was also partially funded by the Wellcome Trust, Medical Research Council, Canadian Institutes of Health Research, Ontario Research Fund. The Nottingham Genetics Group is supported by ARUK and The Big Lottery Fund. The effort from Columbia University was supported by the Taub Institute, the Panasci Fund, the Parkinson's Disease Foundation, and NIH grants NS060113 (L. Clark), P50AG008702 (P.I. Scott Small), P50NS038370 (P.I. R. Burke), and UL1TR000040 (P.I. H. Ginsberg).

O.A.R. is supported by the Michael J. Fox Foundation, NINDS R01# NS078086. The Mayo Clinic Jacksonville is a Morris K. Udall Parkinson's Disease Research Center of Excellence (NINDS P50 #NS072187) and is supported by The Little Family Foundation and by the Mangurian Foundation Program for Lewy Body Dementia research and the Alzheimer Disease Research Center (P50 AG016547). The work from the Mayo Clinic Rochester is supported by the National Institute on Aging (P50 AG016574 and U01 AG006786). This work has received support from The Queen Square Brain Bank at the UCL Institute of Neurology; where TL is funded by an ARUK senior fellowship. Some of the tissue samples studied were provided by the MRC London Neurodegenerative Diseases Brain Bank and the Brains for Dementia Research project (funded by Alzheimer's Society and ARUK). This research was supported in part by both the NIHR UCLH Biomedical Research Centre and the Queen Square Dementia Biomedical Research Unit. This work was supported in part by the Intramural Research Program of the National Institute on Aging, National Institutes of Health, Department of Health and Human Services; project AG000951-12. The University of Pennsylvania case collection is funded by the Penn Alzheimer's Disease Core Center (AG10124) and the Penn Morris K. Udall Parkinson's Disease Research Center (NS053488). The authors would like to thank the Exome Aggregation Consortium and the groups that provided exome variant data for comparison. A full list of contributing groups can be found at http://exac.broadinstitute.org/about. Tissue samples from UCSD are supported by NIH grant AG05131. The authors thank the brain bank GIE NeuroCEB, the French program "Investissements d'avenir" (ANR-10-IAIHU-06). PJT and LM are supported by the Helsinki University Central Hospital, the Folkhälsan Research Foundation and the Finnish Academy.
